## Supplementary figures for "Identification of genomic enhancers through spatial integration of single-cell transcriptomics and epigenomics"

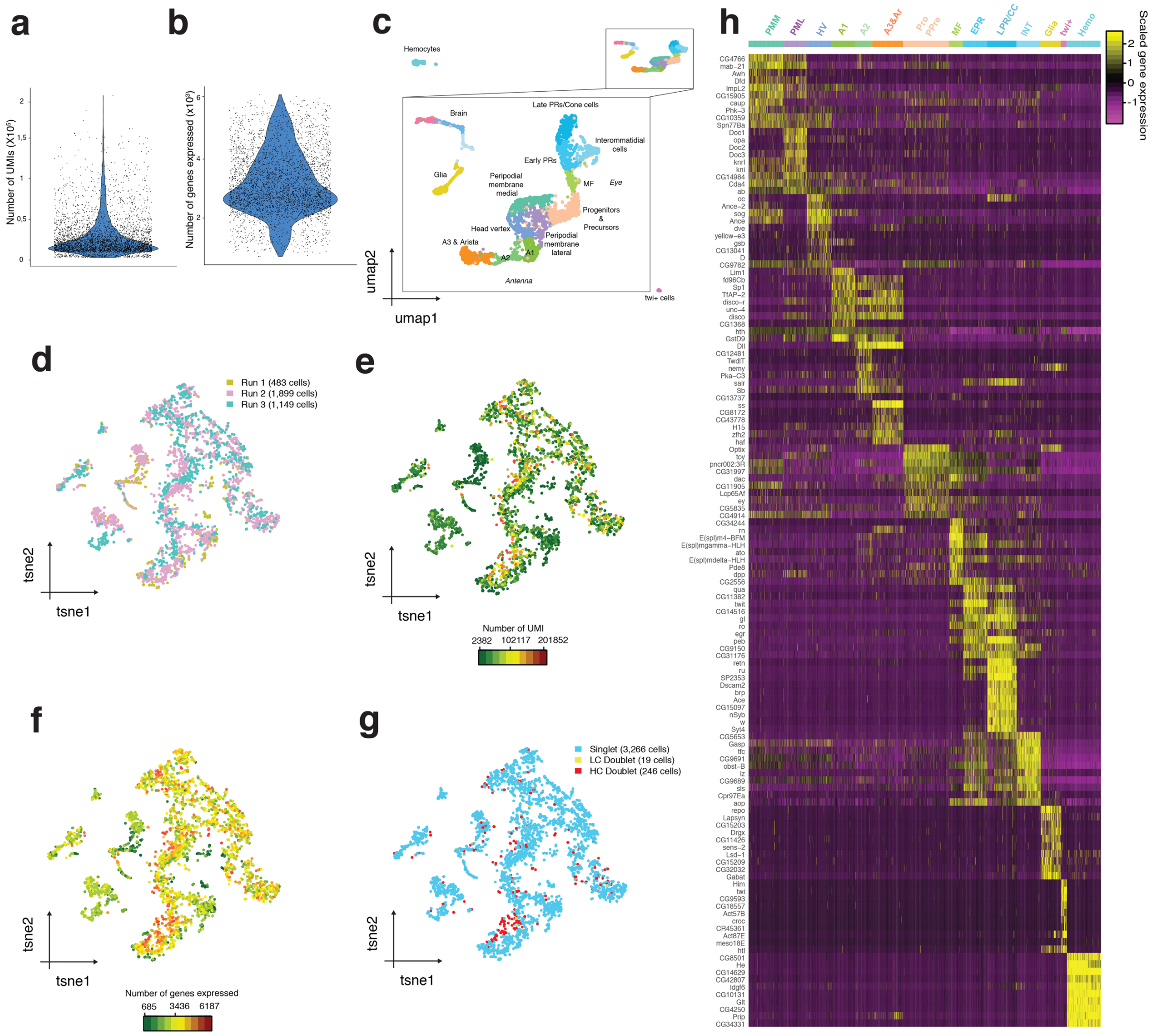


**Figure S1. Quality control and marker genes from the scRNA-seq data set (3,531 cells). a.** Number of UMIs (Unique Molecular Identifiers) per cell. **b.** Number of genes expressed per cell. **c.** Seurat UMAP (3,531 cells) colored by cell type, with a zooming in for the cell types distinct of hemocytes and twi$^{+}$ cells. **d.** Seurat tSNE colored by experimental run. **e.** Seurat tSNE (3,531 cells) colored by the number of UMIs per cell. **f.** Seurat tSNE (3,531 cells) colored by the number of UMIs per cell. **g.** Seurat tSNE colored by singlets, low confidence (LC) doublets and high confidence (HC) doublets as defined by DoubletFinder[1]. **h.** Heatmap showing the scaled expression of the top 10 Seurat markers based on differential expression adjusted p-value per group per cell.

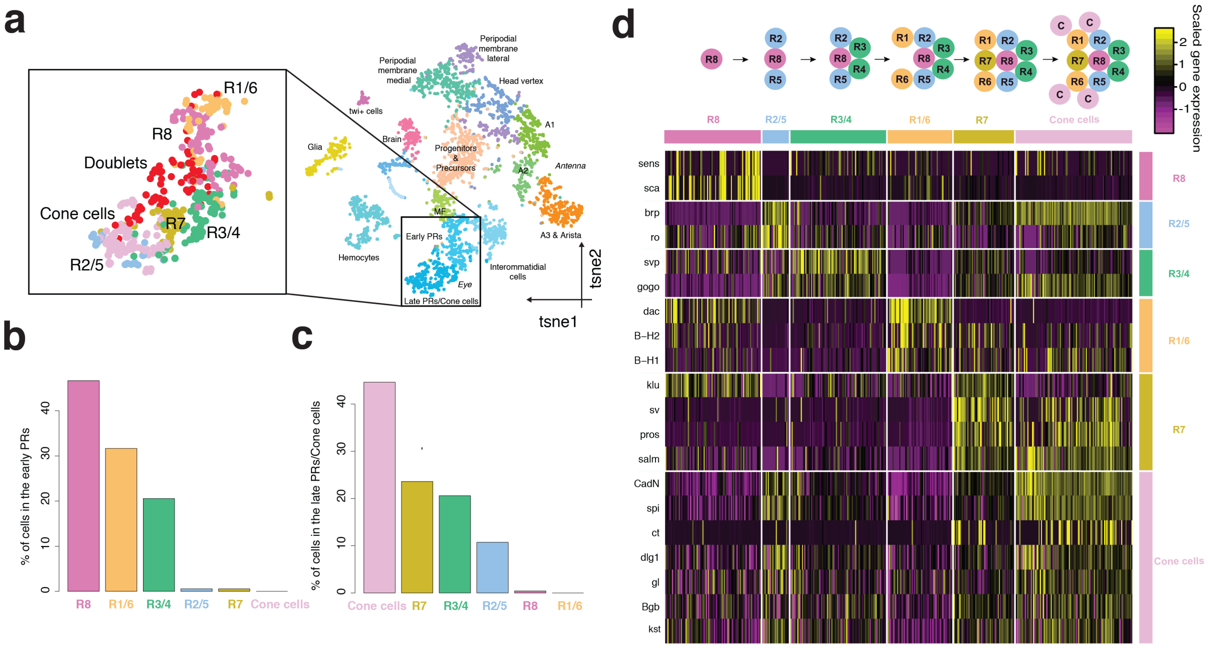


**Figure S2. Identification of photoreceptor subclasses and cone cells in the ommatidial cells. a.** Seurat tSNE annotated by cell type, showing the interommatidial cell clusters annotated by subpopulations augmented. **b.** Percentage of each of the subpopulations in the cluster labeled as early PRs (photoreceptors). **c.** Percentage of each of the subpopulations in the cluster labeled as late PRs/Cone cells. **d.** Heatmap showing known marker genes for each ommatidial subclass. The ommatidial formation process is shown over the heatmap.

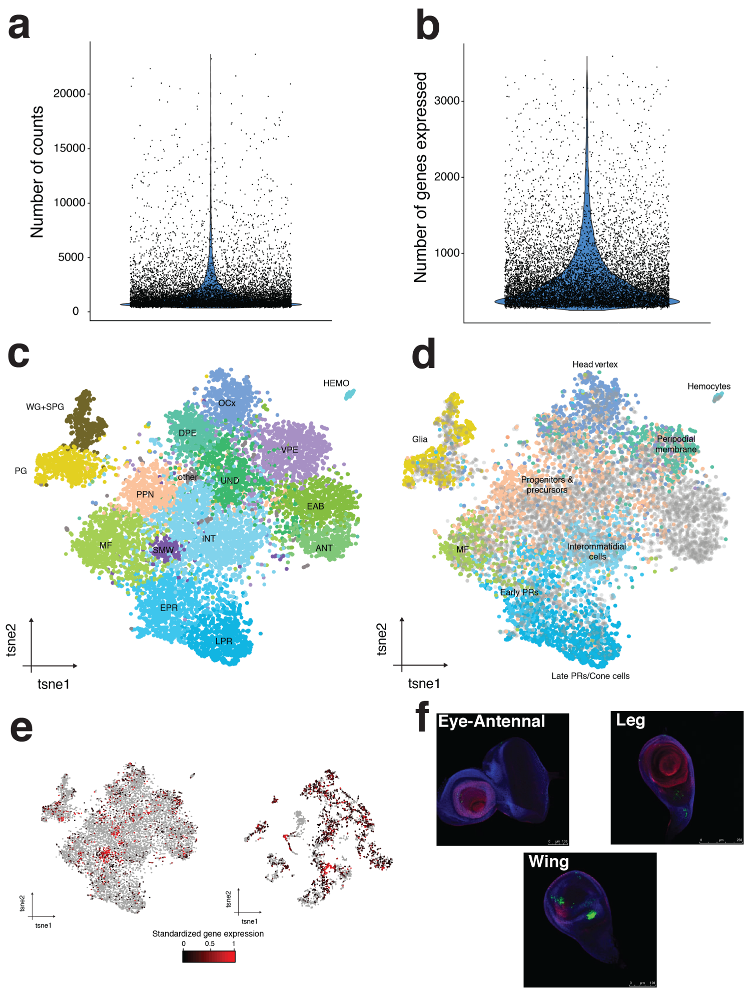


**Figure S3. Comparison with a public Drop-seq data set from the eye disc with 11,500 single cell profiles. a.** Number of gene counts per cell. **b.** Number of genes expressed per cell. **c.** Seurat tSNE annotated by cell type, using the tSNE coordinates provided by Ariss *et al.*[2] WG: Wrappinh glia. SPG: Subperimeural glia. PG: Perinueral glia. HEMO: Hemocytes. OCx: Ocellar complex. DPE: Dorsal peripodial membrane. VPE: Ventral peripodial membrane. EAB: Eye-antennal border. ANT: Antenna. UND: Undifferentiated cells. PPN: Pre-proneural domain. MF: Morphogenetic furrow. SMW: Second mitotic wave. INT: Interommatidial cells. EPR: Early photoreceptors. LPR: Late Photoreceptors. **d.** Seurat tSNE from Ariss et al., colored by label transfer from our data set. Cells in grey were removed from the analysis due to low depth. **e.** Expression of Claspin, marker gene of the second mitotic wave cells, in the Drop-seq tSNE (left) and our data set (right, with 3,531 cells) **f.** Images retrieved from Janelia Flylight[3] showing the activity of the enhancer linked to twi, marker of adepithelial cells, in the eye-antennal disc, the leg disc and the wing disc.

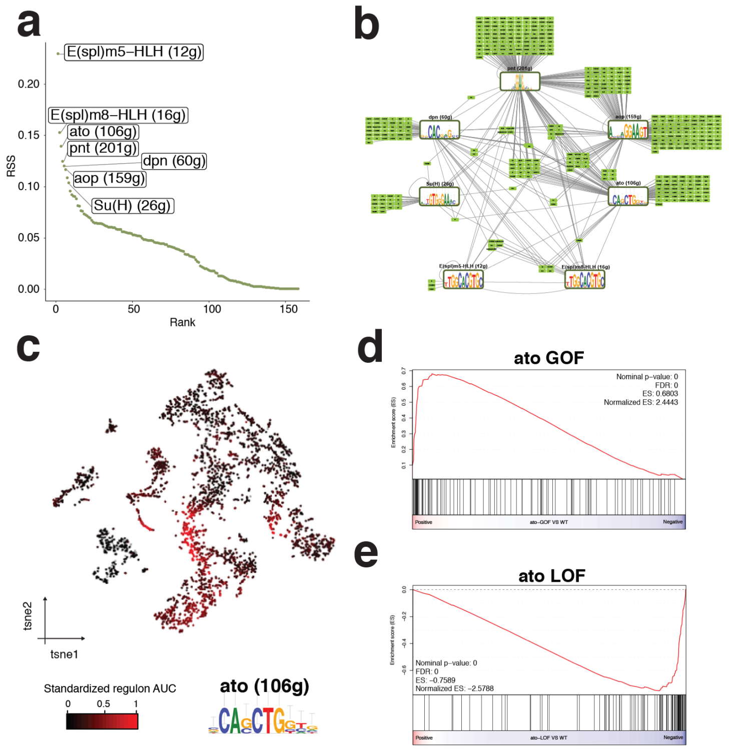


**Figure S4. Validation of the gene regulatory networks in the morphogenetic furrow. a.** Regulon Specifity Score (RSS) for the morphogenetic furrow. The top 7 regulons are highlighted. **b.** Gene Regulatory Network of the morphogenetic furrow based on the top 7 regulons. Transcription factors and their target genes are shown. **c.** Seurat tSNE (3,531 cells) coloured by the enrichment of the atonal regulon. The top enriched atonal motif on the regulon is shown. **d.** GSEA plot showing the enrichment of the predicted atonal targets on an atonal gain-of-function (GOF) mutant. The gene ranking is based on the logFC from the differential gene expression analysis of the atonal gain-of-function mutant versus wild type[4]. **e.** GSEA plot showing the enrichment of the predicted atonal targets in an atonal loss-of-function (LOF) mutant. The gene ranking is based on the logFC from the differential gene expression analysis of the atonal loss-of-function mutant versus wild type[4].

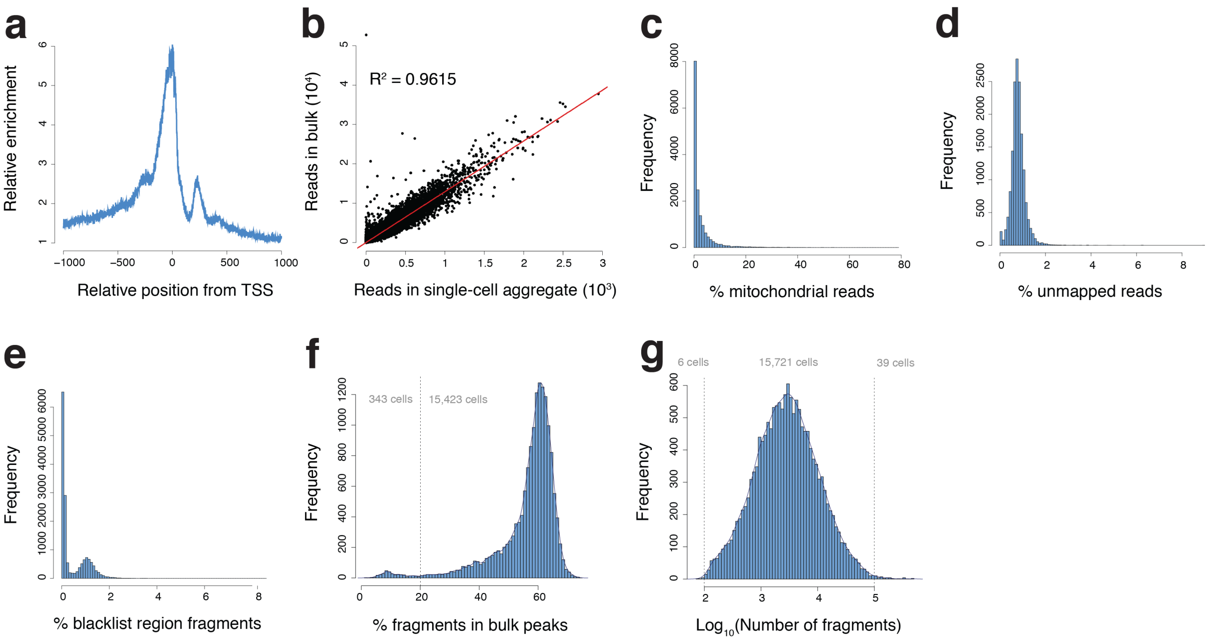


**Figure S5. Quality control of scATAC-seq data. a.** Relative enrichment of the accessibility signal versus position from the TSS. TSS has higher relative accessibility compared to surrounding areas. **b.** Correlation between the accessibility of regions in the bulk ATAC-seq and the aggregated single-cell profiles. **c.** Percentage of mitochondrial reads. **d.** Percentage of unmapped reads. **e.** Percentage of fragments in blacklisted regions. **f.** Percentage of fragments in bulk peaks. Cells with less than 20% of the fragments in bulk peaks, were filtered out. **g.** Normalized ($log_{10}$) number of fragments. Cells with less than 100 fragments or more than 100,000 fragments were filtered out.

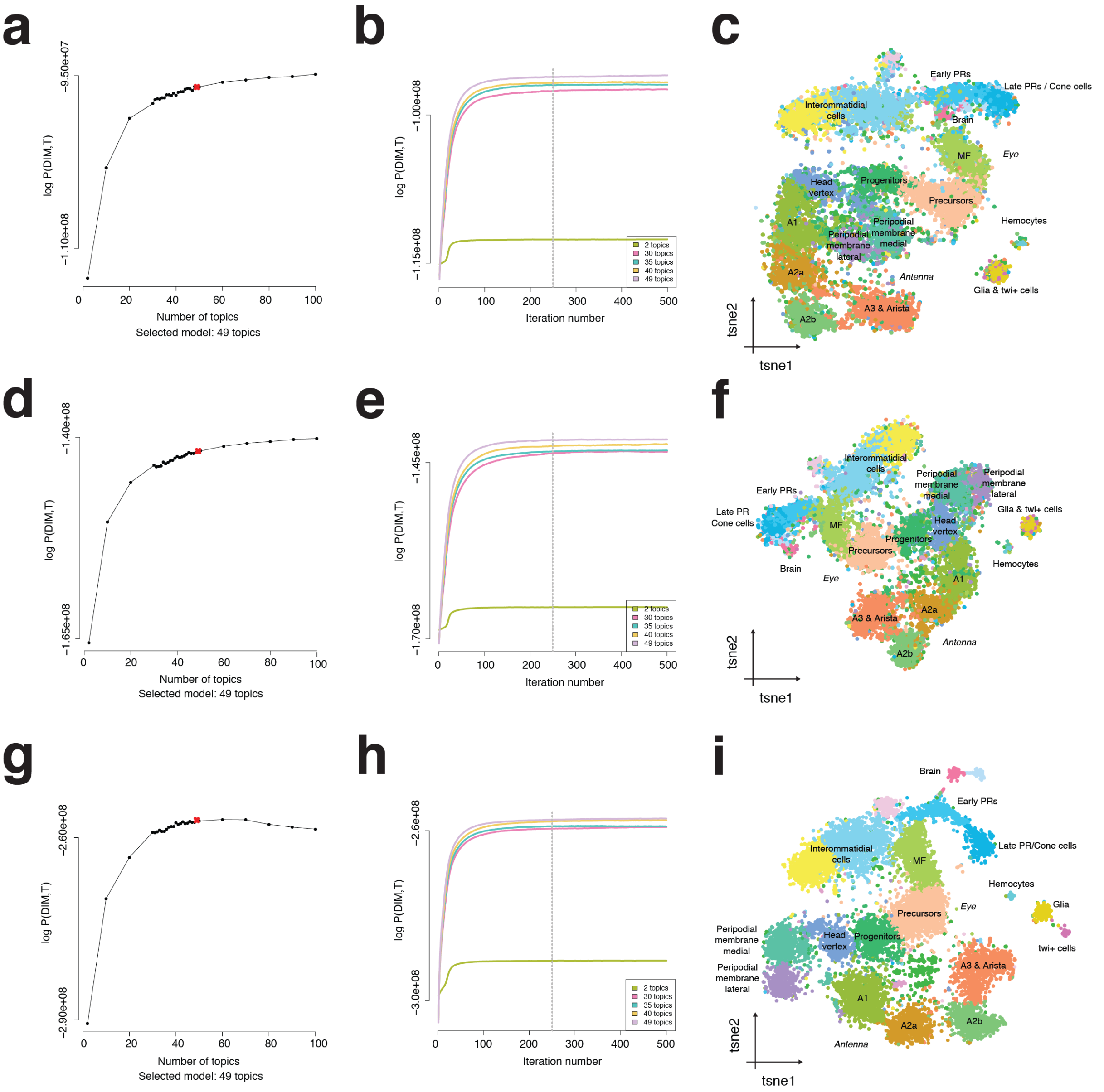


**Figure S6. Selection of regulatory regions for the analysis of scATAC-seq data. a-c.** cisTopic modelling using bulk narrow peaks as called by MACS as regulatory regions (13,567 regions). The log-likelihood per model, the log-likelihood per iteration for selected models, and the cisTopic cell tSNE (with 15, 387 cells) of the selected model (with 49 topics) are shown. **d-f.** cisTopic modelling using the summits as called by MACS extended +/- 250 bp as regulatory regions (16,417 regions). The log-likelihood per model, the log-likelihood per iteration for selected models, and the cisTopic cell tSNE of the selected model (with 49 topics) are shown. **g-h.** cisTopic modelling using the cisTarget regions as regulatory regions (129,553 regions). The log-likelihood per model, the log-likelihood per iteration for selected models, and the cisTopic cell tSNE of the selected model (with 49 topics) are shown.

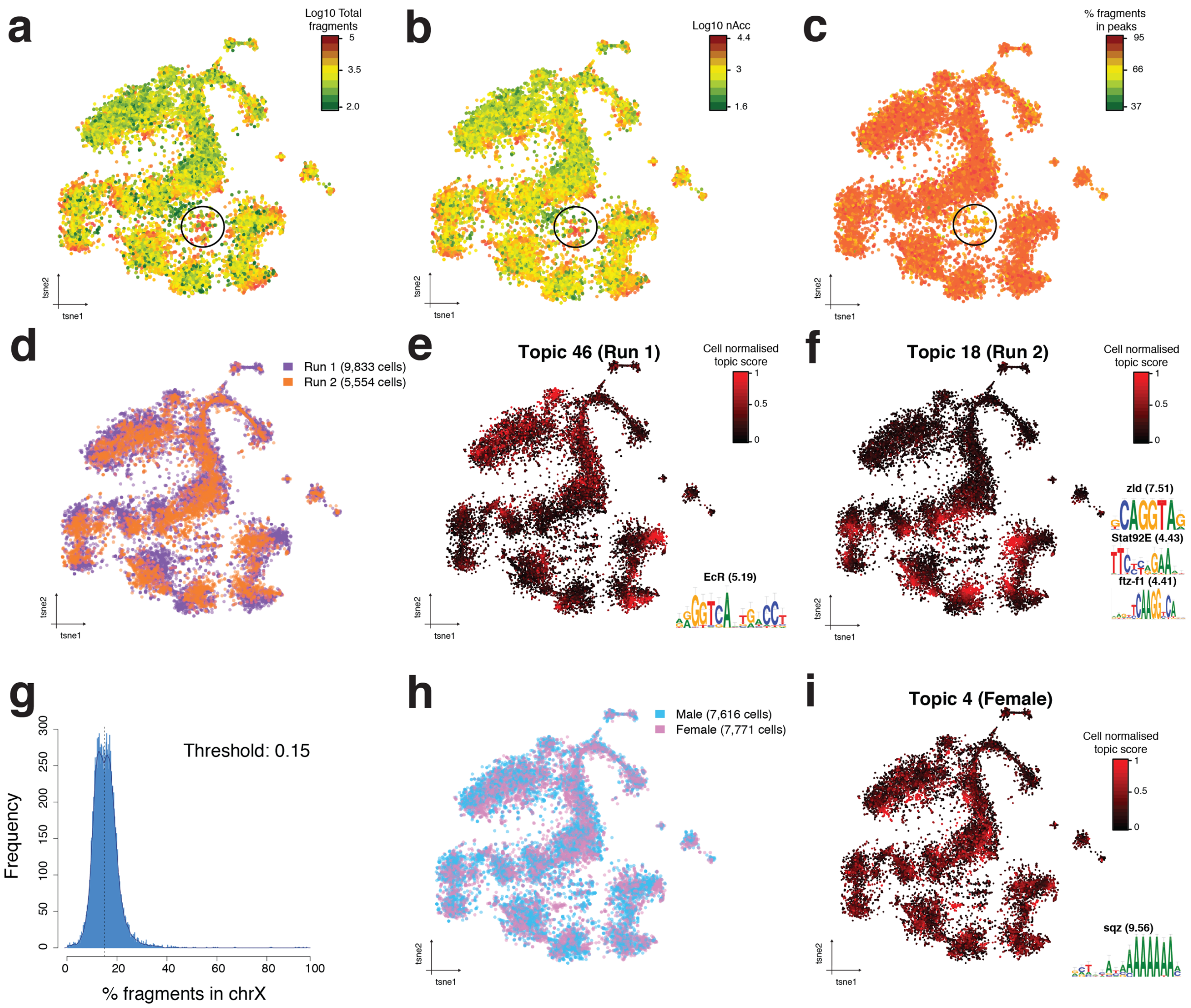


**Figure S7. Cell clustering is driven by cell type and not by batch effects. a.** cisTopic cell tSNE (15,387 cells) colored by normalized number of fragments ($log_{10}$). **b.** cisTopic cell tSNE colored by normalized number of accessible regions ($log_{10}$) **c.** cisTopic cell tSNE colored by percentage of fragments that overlap bulk peaks. **d.** cisTopic cell tSNE colored by experimental run. **e.** cisTopic cell tSNE colored by topic 46 enrichment. A representative enriched motif with Normalised Enrichment Score (NES) is shown. **f.** cisTopic cell tSNE colored by topic 18 enrichment. Representative enriched motifs with NES are shown. **g.** Histogram showing the bimodal distribution of fragments over the X chromosome. Cells with more than 15% of fragments on the X chromosome are classified as females; otherwise, as males. **h.** cisTopic cell tSNE colored by assigned sex. **i.** cisTopic cell tSNE colored by topic 4 enrichment. A representative enriched motif with NES is shown.

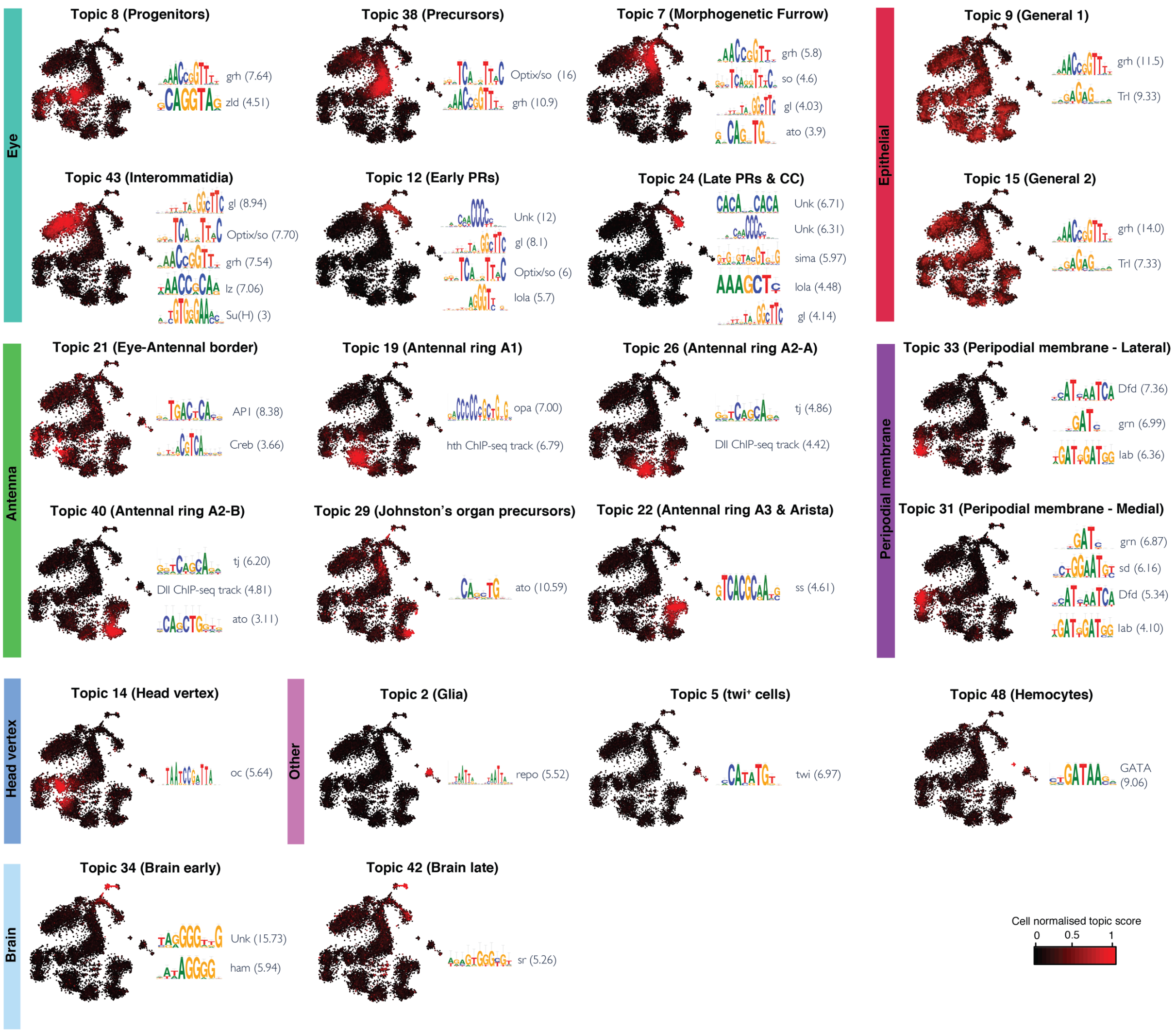


**Figure S8. Motif enrichment on the regions within regulatory topics reveal key master regulators of the cell populations in the eye-antennal disc.** For each selected topic, cisTopic cell tSNE colored by topic enrichment accompanied by representative enriched motifs (which are linked to known master regulators of each cell type) with Normalized Enrichment Score (NES).

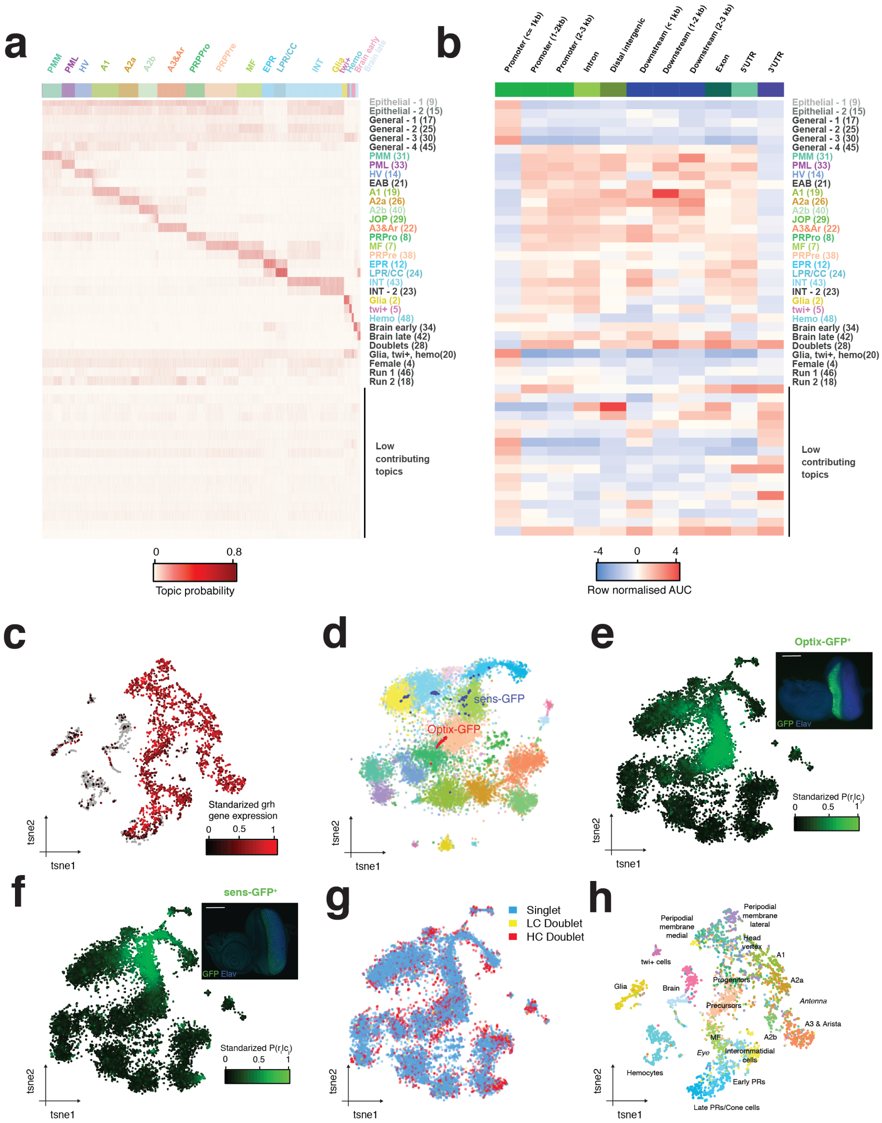

**Figure S9. Annotation and validation of topics and cell clusters from scATAC-seq profiles. a.** cisTopic cell-topic heatmap showing the probability of each topic in each cell and the topic annotation. **b.** Heatmap showing the enrichment of each regions class in each topic. **c.** Seurat scRNA-seq tSNE colored by grh gene expression. **d.** cisTopic cell tSNE after the integration of FAC-sorted cells profiled by Fluidigm C1. Cells profiled with 10X are shown with transparency colored by annotation. **e.** cisTopic cell tSNE colored by the probability of the region corresponding to the Optix2/3 enhancer, whose activity is measured by GFP signal in the adjacent image. Scale bar: 100um. **f.** cisTopic cell tSNE colored by the probability of the region corresponding to the sens-F2 enhancer, whose activity is measured by GFP signal in the adjacent image. Scale bar: 100um. **g.** Annotation of singlets and doublets based on DoubletFinder (on the gene accessibility matrix). **h.** Seurat scRNA-seq tSNE colored by the labels transferred from the scATAC-seq annotation. PMM: Peripodial Membrane Medial. PML: Peripodial Membrane Lateral. HV: Head Vertex. Pro: Progenitors. Pre: Precursors. MF: Morphogenetic Furrow. EPR: Early photoreceptors. LPR/CC: Late photoreceptors and cone cells. INT: Interommatidial cells. Hemo: Hemocytes. EAB: Eye-antennal border. JOP: Johnston Organ Precursor.

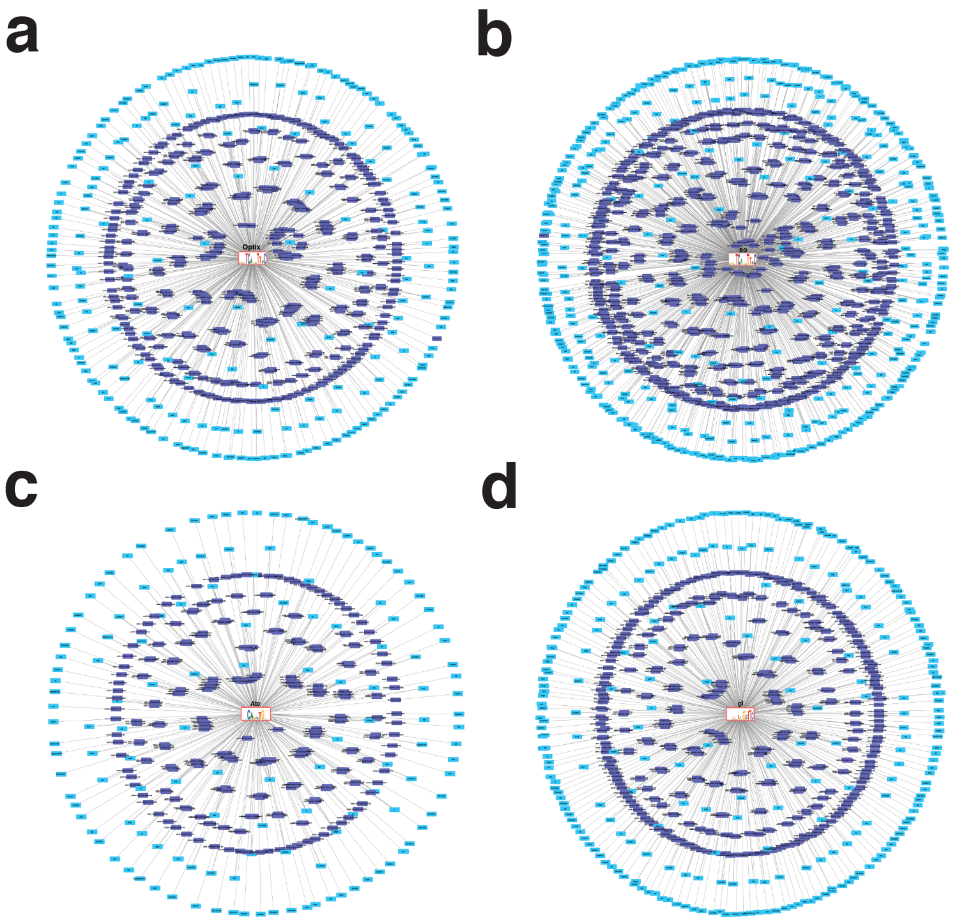


**Figure S10. Gene Regulatory Networks based on co-expression from the scRNA-seq data and enhancer accessibility and motif enrichment from the scATAC-seq data.** Gene Regulatory Networks containing master TF (with motif, in red), target enhancer regions (in purple), and target genes (in blue), which are co-expressed with the TF in the scRNA-seq data and have a target enhancer in their surroundings (within +- 5kb from the TSS and introns). Examples are shown for Optix (a), so (b), ato (c) and gl (d).

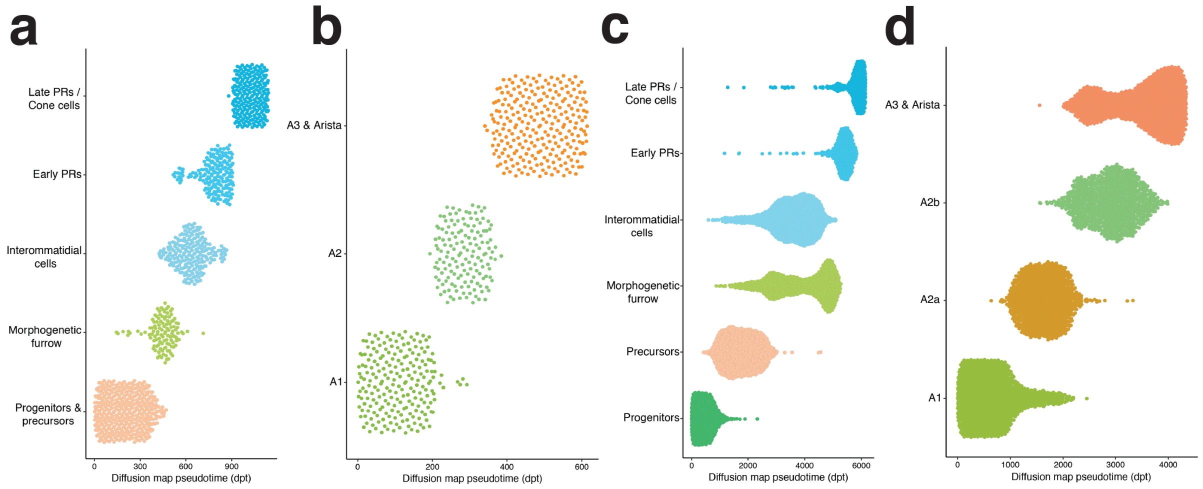


**Figure S11. Ordering cell types in the eye and antenna by pseudotime. a.** Eye cell types ordered by pseudotime using scRNA-seq data. **b**. Antennal cell types ordered by pseudotime using scRNA-seq data. **c.** Eye cell types ordered by pseudotime using scATAC-seq data. **d.** Antennal cell types ordered by pseudotime using scATAC-seq data.

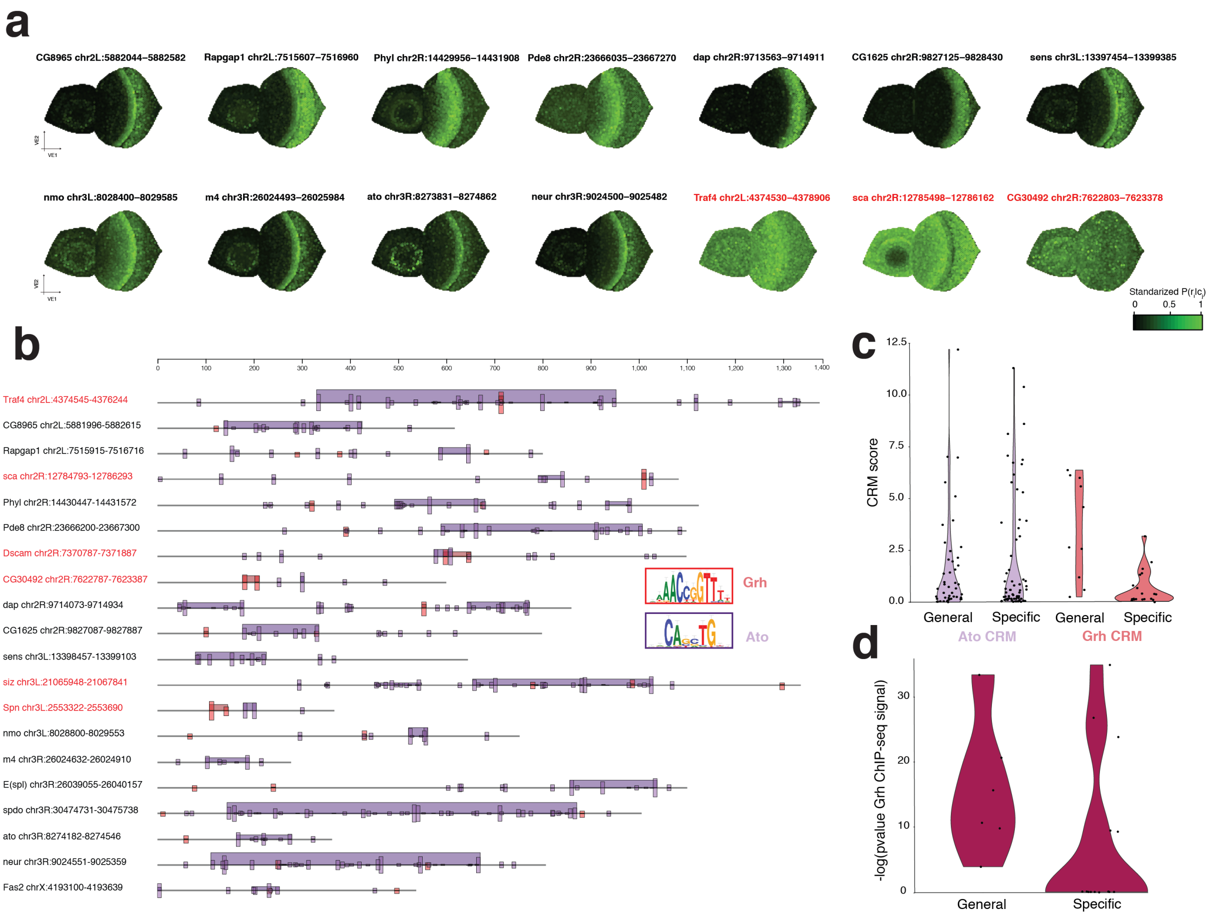


**Figure S12. Uncoupling enhancer accessibility and activity in validated atonal enhancers from Aerts *et al.* a.** Virtual eye-antennal disc colored by the accessibility probability of 14 of the atonal enhancers. Enhancers with general accessibility and enriched for grh binding sites are shown in red. **b.** TOUCAN view showing the location of atonal (purple) and grh (red) motifs and Cis-Regulatory Modules (CRMs). Enhancers with general accessibility and enriched for grh binding sites are shown in red. **c.** Violin plot comparing CRMs scores in specific and generally accessible regions. **d.** Violin plot comparing MACS2 log10(p-val) of the grh ChIP-seq peaks in specific and generally accessible regions. Jacobs *et al*.[5] showed that Grh and Ato enhancers had a higher ATAC-seq signal compared to uniquely Ato targets. Here we find that these differences in signal are due to Ato enhancers being uniquely accessible in the morphogenetic furrow, while shared Grh and Ato enhancers are accessible across all epithelial cells.

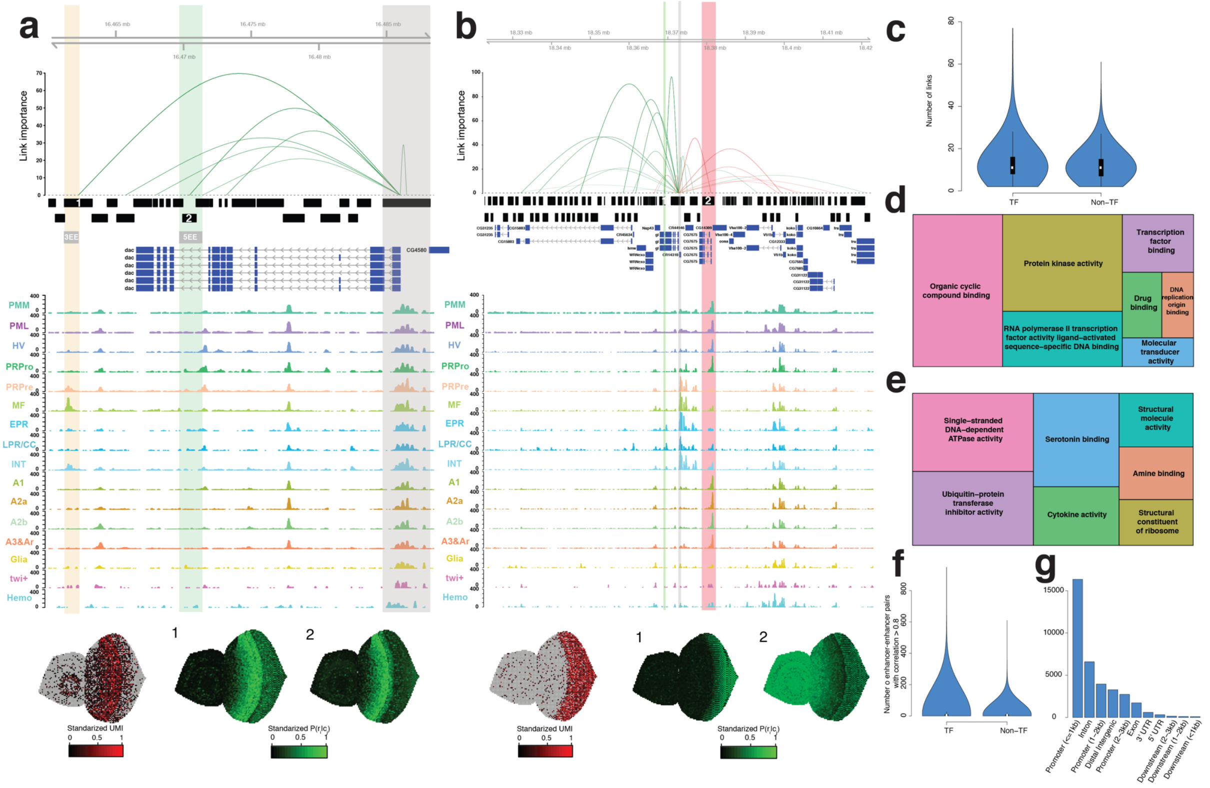


**Figure S13. Enhancer-to-target links support basic principles of gene regulation. a.** Enhancer-to-target links for the dac gene. From top to bottom are shown: position, link importance, cisTarget regions, validated dac enhancers, gene annotation and scATAC-seq aggregates by cell type. The expression of dac is shown on the virtual eye in a red scale, the accessibility probability of 3EE (1) and 5EE (2) is shown in a green scale. 3EE is highlighted in orange, 5EE in green and the promoter of dac in grey. **b.** Enhancer-to-target links for the gl gene. From top to bottom are shown: position, link importance, cisTarget regions, gene annotation and scATAC-seq aggregates by cell type. The expression of gl is shown on the virtual eye in a red scale, the accessibility probability of the top activating region based on importance (highlighted in green, 1) and top repressive region (highlighted in red, 2) are shown in green. Gl promoter is highlighted in grey. **c.** Number of links per TF and non-TF gene. **d.** Revigo view of the functional GO terms enriched found by GOrilla in the genes with a high number of links (based on the decreasing ranking by number of genes of the links). **e.** Revigo view of the functional GO terms enriched found by GOrilla in the genes with a low number of links (based on the ascending ranking by number of genes of the links). **f.** Number of enhancer-enhancer pairs with a correlation above 0.8 (based on the accessibility probability) for TF and non-TF genes. **g.** Classification of the regions involved in the links. PMM: Peripodial Membrane Medial. PML: Peripodial Membrane Lateral. HV: Head Vertex.

Pro: Progenitors. Pre: Precursors. MF: Morphogenetic Furrow. EPR: Early photoreceptors. LPR/CC: Late photoreceptors and cone cells. INT: Interommatidial cells. Hemo: Hemocytes. JOP: Johnston Organ Precursor.

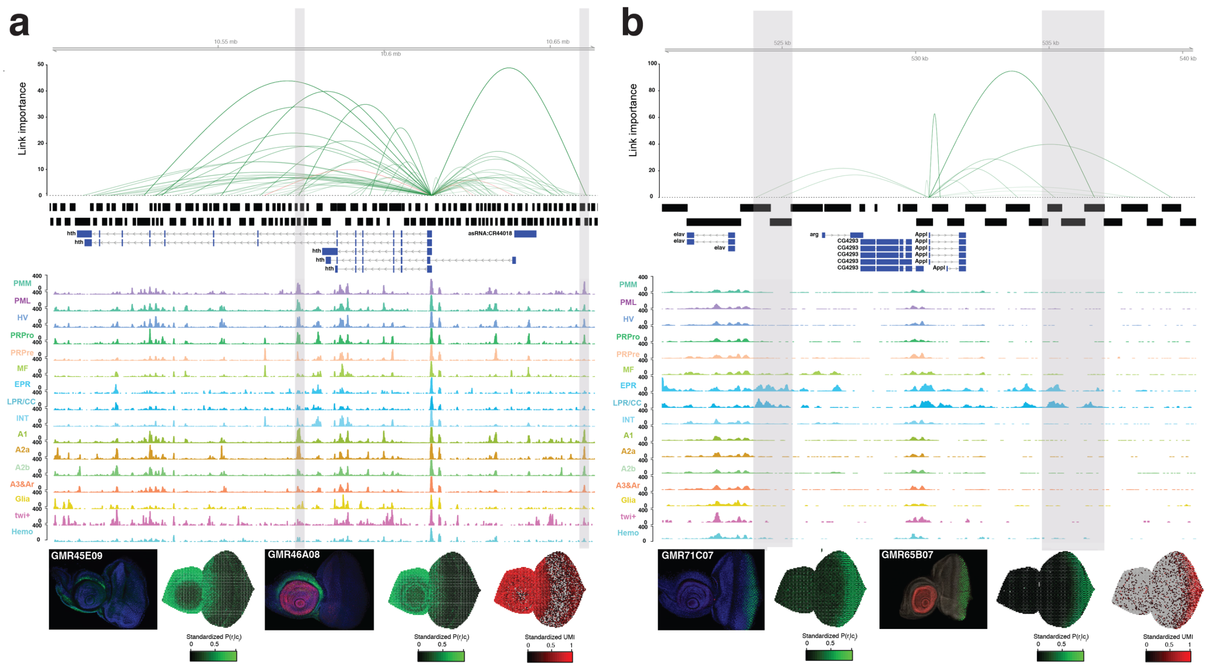


**Figure S14. Genes are regulated by many enhancers, some of which show redundant activity (and accessibility).** Enhancer-to-target links for hth (a) and Appl. Top: Links between enhancer and target genes, whose height represent their importance. Midde top: cisTarget regions. Middle: Genome annotation. Middle bottom: Normalized aggregate ATAC-seq profiles per cell type. Bottom: Enhancer activity (images taken from the Janelia Flylight Project), and virtual eye colored by the predicted enhancer accessibility (green) and the expression of the gene (red). Redundant enhancers are highlighted in grey.

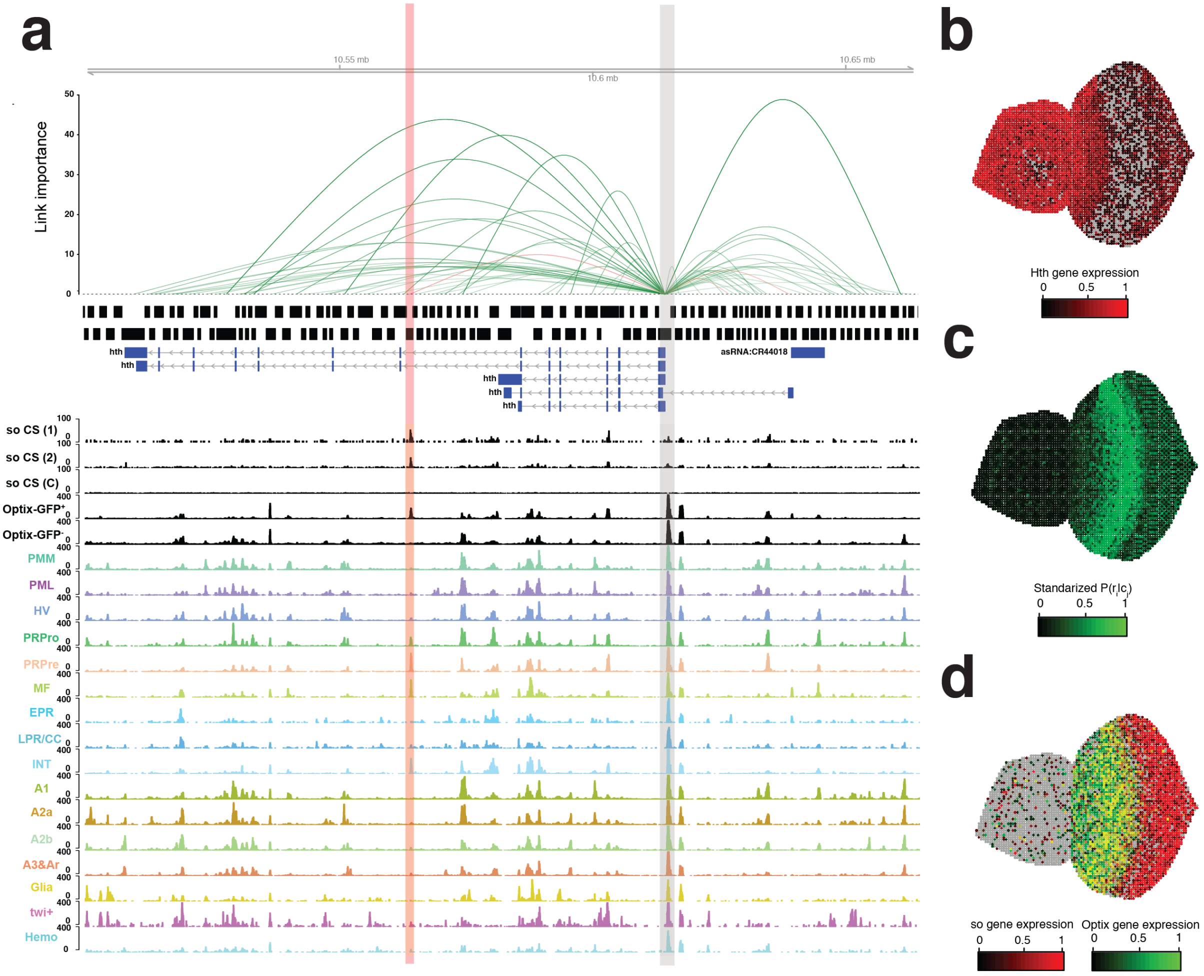


**Figure S15. Hth is potentially repressed by an enhancer bound by so and also accessible in FAC-sorted Optix-GFP$^+$ cells. a.** Enhancer-to-target links for the hth gene. From top to bottom are shown: position, link importance, cisTarget regions, gene annotation, so ChIP-seq data (replicate 1, replicate 2 and control), Optix-GFP$^+$ cells bulk ATAC-seq profile, Optix-GFP$^-$ cells bulk ATAC-seq profile and scATAC-seq aggregates by cell type. The repressive enhancer is highlighted in red, the promoter of hth is highlighted in grey. **b.** Virtual eye-antennal disc colored by the standarized gene expression of hth. **c.** Virtual eye-antennal disc colored by the standarized accessibility probability of the represive enhancer. **d.** Virtual eye-antennal disc colored by the standarized gene expression of so (red) and Optix (green). PMM: Peripodial Membrane Medial. PML: Peripodial Membrane Lateral. HV: Head Vertex. Pro: Progenitors. Pre: Precursors. MF: Morphogenetic Furrow. EPR: Early photoreceptors. LPR/CC: Late photoreceptors and cone cells. INT: Interommatidial cells. Hemo: Hemocytes. JOP: Johnston Organ Precursor.

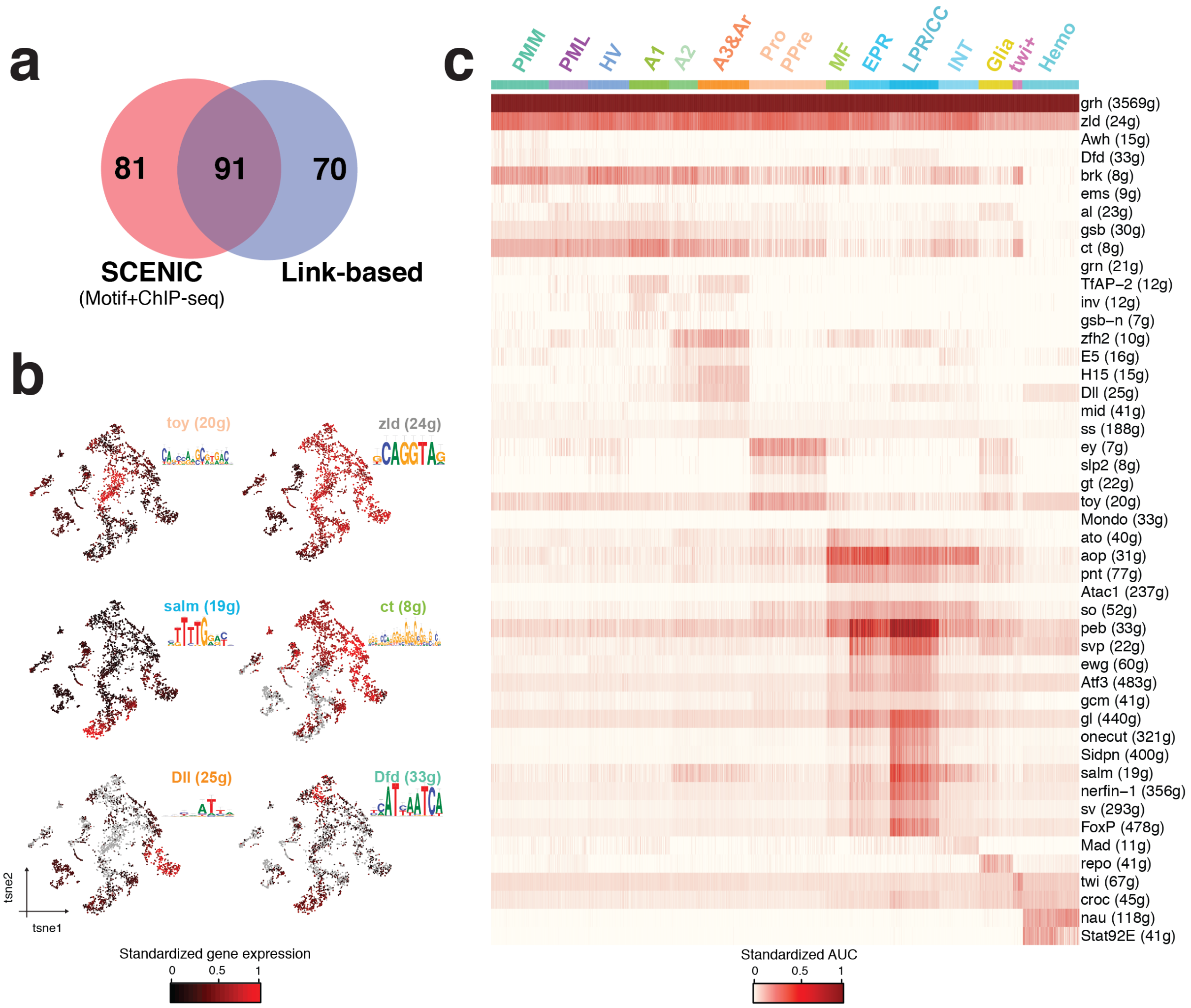


**Figure S16. Link-based regulons identify additional master regulators during the eye-antennal disc development. a.** Venn diagram showing the overlap between SCENIC and link-based regulons. **b.** Seurat scRNA-seq tSNE colored by regulon enrichment in each cell. **c.** Cell-to-regulon heatmap showing the standardized enrichment or Area Under the Curve (AUC) from SCENIC for each selected regulon based on RSS in each cell. PMM: Peripodial Membrane Medial. PML: Peripodial Membrane Lateral. HV: Head Vertex. Pro: Progenitors. Pre: Precursors. MF: Morphogenetic Furrow. EPR: Early photoreceptors. LPR/CC: Late photoreceptors and cone cells. INT: Interommatidial cells. Hemo: Hemocytes. JOP: Johnston Organ Precursor.

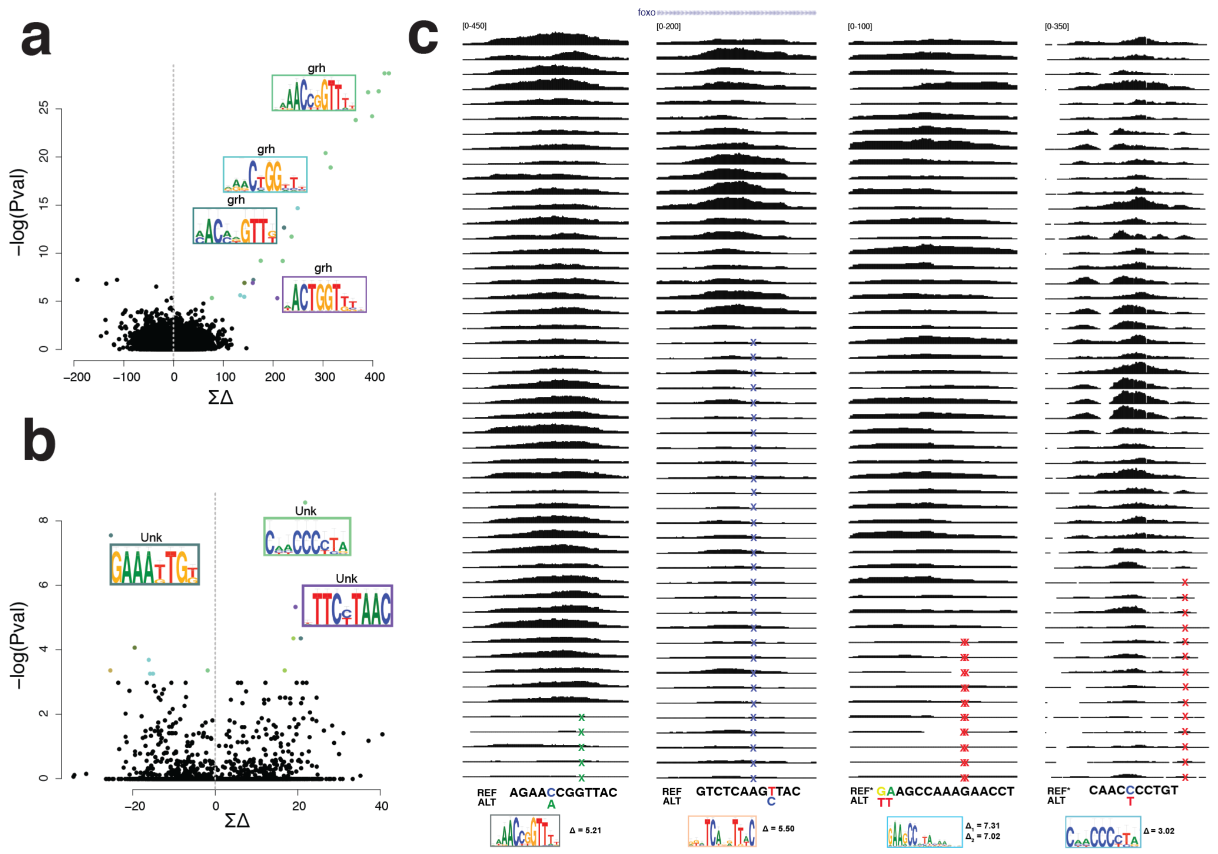


**Figure S17. Cell-type specific analysis of chromatin accessibility QTLs reveals additional motifs involved chromatin opening compared to bulk analysis. a.** Delta motif plot, showing the cumulative effect of the 10,969 caQTLs found genome-wide on the CRM score of 24,454 motifs on the x-axis. The y-axis shows the enrichment of motifs affected by caQTLs compared to control SNPs genome-wide (one-sided Fisher's exact test, log transformed). **b.** Delta motif plot, showing the cumulative effect of the 323 caQTLs found on regions specifically accessible in late PRs and cone cells (Topic 24) on the CRM score of 24,454 motifs on the x-axis. The y-axis shows the enrichment of motifs affected by caQTLs compared to controls SNPs (one-sided Fisher's exact test, log transformed). **c.** Bulk ATAC-seq profiles of the inbred lines, on regions affected by caQTLs that modify the highlighted motifs. The caQTLs coordinates are, from left to right: chr3L:17392596, chr3R:14076593, chr2R:18674001 and chr2R:18674002, and chr3R:29376820. REF sequences with an * indicate that the reverse complement strand is given.

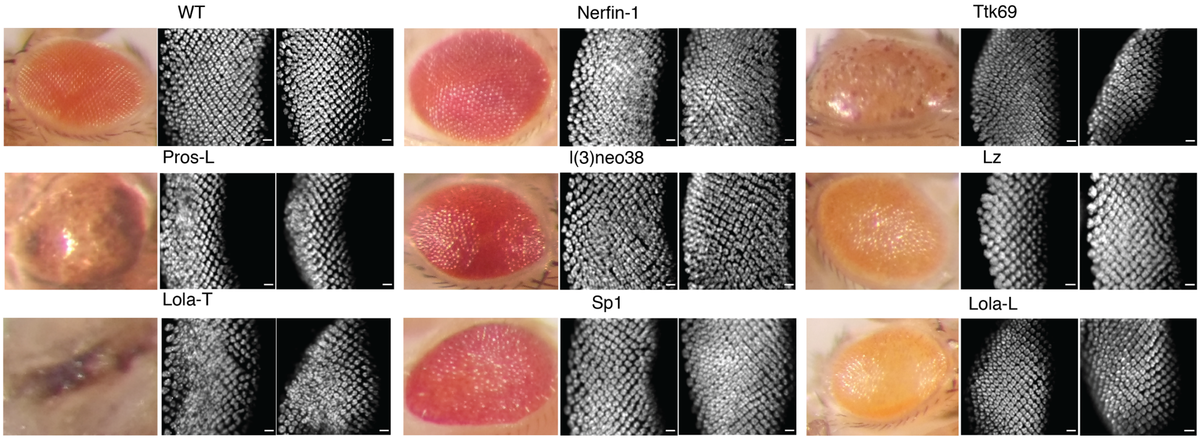


**Figure S18. Overexpression of 13 TFs using the GMR-GAL4/UAS-TF system reveals 8 master regulators whose overexpression causes defects during the development of the eye disc.** The adult eye in each of these lines is shown (except for Pros and Lola-T), together with the elav staining on posterior to the morphogenetic furrow in the third instar larvae eye disc on two biological replicates. Scale bar: 10 um.

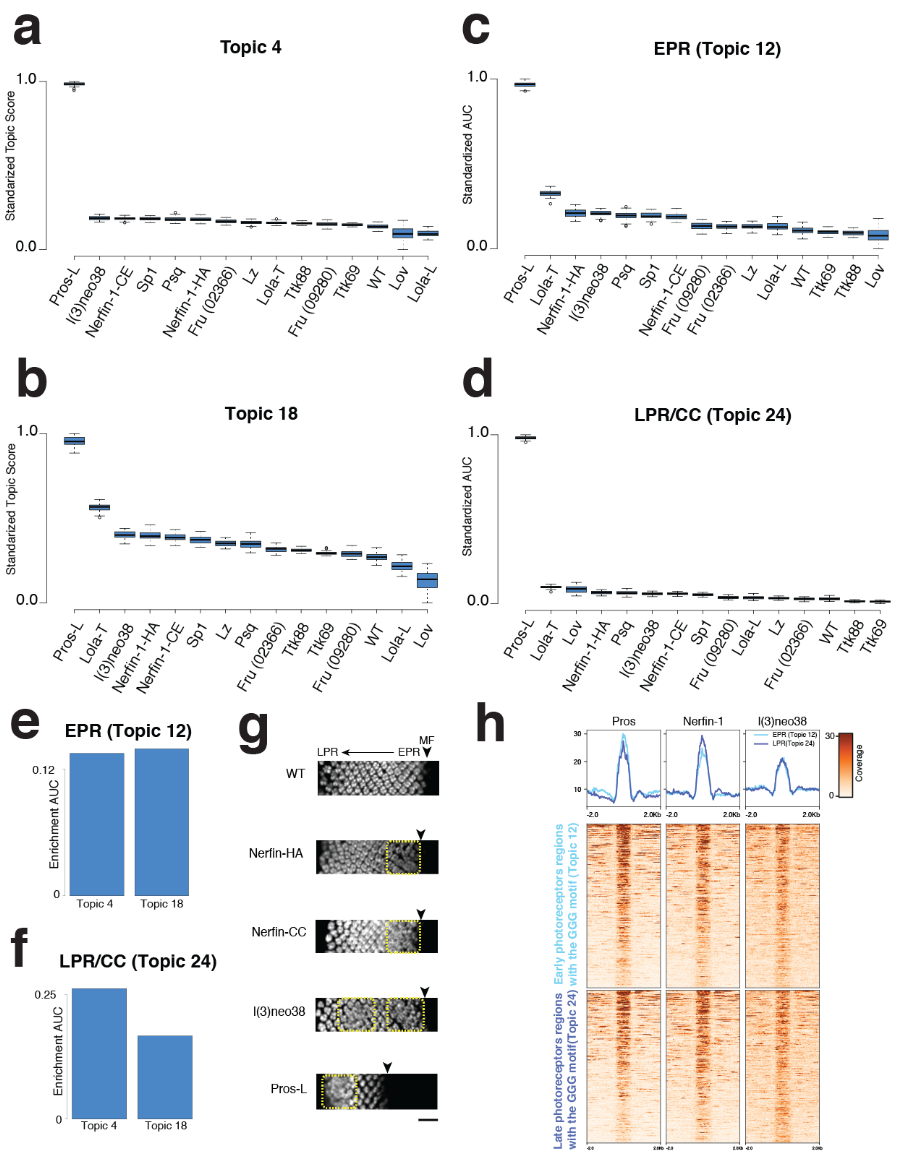


**Figure S19. Pros-L overexpression result on the opening of GGG enriched regions. a.** Standarized topic 4 enrichment (from the bootstrapped analysis) on the bootstrapped samples. **b.** Standarized topic 18 enrichment (from the bootstrapped analysis) on the bootstrapped samples. **c.** Standarized enrichment of the early photoreceptor regions (topic 12) on the bootstrapped samples. **d.** Standarized enrichment of the late photoreceptor regions (topic 24) on the bootstrapped samples. **e.** AUC enrichment of the early photoreceptor regions on the topics 4 and 18 (from the bootstrapped analysis). **f.** AUC enrichment of the late photoreceptor regions on the topics 4 and 18 (from the bootstrapped analysis). **g.** Elav staining on the third instar larvae eye-antennal disc (posterior to the morphogenetic furrow, marked with an arrow) of the selected GMR-GAL4 UAS-TF and wild type lines. Scale bar: 20 µm.**h.** Heatmaps showing the normalized coverage of early photoreceptor regions (topic 12) enriched in the GGG motif and late photoreceptor regions (topic 24) enriched in the GGG motif on the ChIP-seq profiles of Prospero, Nerfin-1 and l(3)neo38.

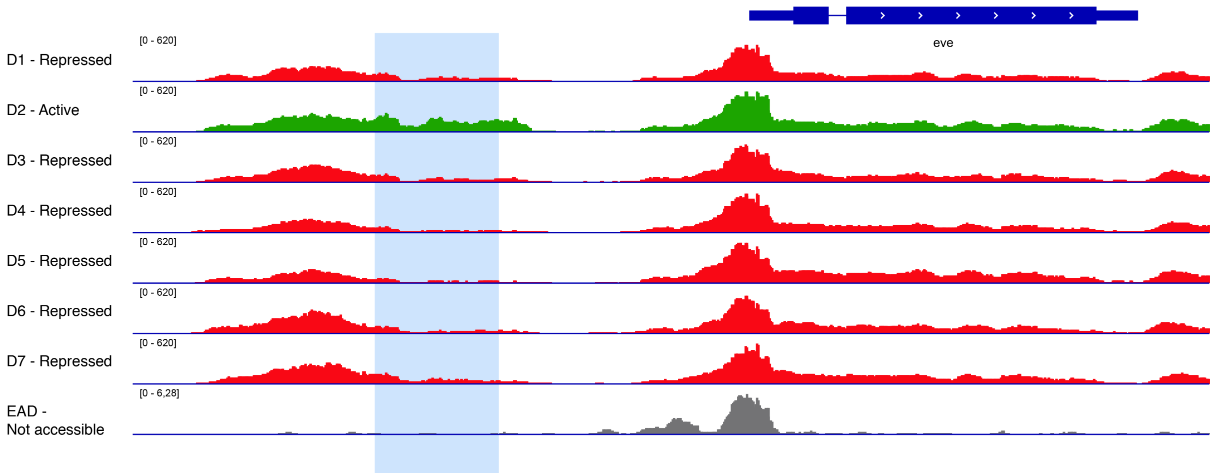


**Figure S20. Repressed enhancers show a reduced ATAC-seq signal.** Normalized ATAC-seq profiles of the different embryonic domains (D1-7, as described by Bozek *et al.*[6]) and a wild type third instar larvae eye antennal disc profile. The eve stripe 2 enhancer is only active in the second embryonic stripe (D2), while repressed in the rest of the embryo. In the eye-antennal disc this enhnacer is neither activated or repressed.
